# CryoSamba: self-supervised deep volumetric denoising for cryo-electron tomography data

**DOI:** 10.1101/2024.07.11.603117

**Authors:** Jose Inacio Costa-Filho, Liam Theveny, Marilina de Sautu, Tom Kirchhausen

## Abstract

Cryogenic electron tomography (cryo-ET) has rapidly advanced as a high-resolution imaging tool for visualizing subcellular structures in 3D with molecular detail. Direct image inspection remains challenging due to inherent low signal-to-noise ratios (SNR). We introduce CryoSamba, a self-supervised deep learning-based model designed for denoising cryo-ET images. CryoSamba enhances single consecutive 2D planes in tomograms by averaging motion-compensated nearby planes through deep learning interpolation, effectively mimicking increased exposure. This approach amplifies coherent signals and reduces high-frequency noise, substantially improving tomogram contrast and SNR. CryoSamba operates on 3D volumes without needing pre-recorded images, synthetic data, labels or annotations, noise models, or paired volumes. CryoSamba suppresses high-frequency information less aggressively than do existing cryo-ET denoising methods, while retaining real information, as shown both by visual inspection and by Fourier shell correlation analysis of icosahedrally symmetric virus particles. Thus, CryoSamba enhances the analytical pipeline for direct 3D tomogram visual interpretation.

## INTRODUCTION

Cryogenic electron tomography (cryo-ET) has become an important tool in structural biology for imaging three-dimensional biological structures with molecular resolution in their native context (Baumeister et al., 1999; Medalia et al., 2002). Achieving this dual capability requires an extremely low electron dose per tilt image to avoid sample damage, resulting in data with very low signal-to-noise ratios (SNR) (Gan et al., 2012). Enhancement of signal through downstream processing can be achieved by aligning and merging multiple instances of an invariant biological structure (when one exists), an approach known as subtomogram averaging (STA) (Wan et al., 2016). STA has effectively produced high-resolution electron density maps of viruses (Schur et al., 2016), ribosomes (Erdmann et al., 2021), and nuclear pores (Mosalaganti et al., 2022).

Traditionally, contrast in cryoET volumes has been enhanced using low pass filtering and pixel binning (Lučić et al., 2005). Recently, deep learning approaches have emerged as superior alternatives (Buchholz et al., 2019; Bepler et al., 2020). These methods adapt to the intricacies of the data, but because cryo-ET data generally lack ground truth high SNR images for direct supervision, most deep-learning denoising algorithms rely on self-supervision (Lehtinen et al., 2018), using paired 3D volumes from an even/odd split of the cryo-ET tilt-series (Buchholz et al., 2019; Bepler et al., 2020), synthetic or annotated data (Zeng et al., 2024), or noise modeling (Li et al., 2022).

Despite their effectiveness in enhancing SNR, these approaches inevitably distort the original data, particularly by suppressing high-frequency details (Bepler et al., 2020). Consequently, denoised tomograms are typically reserved for interpretability tasks, such as identifying the regions of interest for STA or other downstream processing, which then uses the original raw data. These distortions can impede interpretability if high spatial frequency details are essential for distinguishing objects of interest. Therefore, it is desirable for cryo-ET pipelines to incorporate denoising methods that minimize such deformations.

We have developed a deep learning-based denoising method for cryo-ET that enhances contrast with minimal deformation when compared to other current techniques. CryoSamba, the software that applies this approach, operates in a fully self-supervised manner, training directly on the raw three-dimensional volume without requiring additional data such as paired volumes or simulations. CryoSamba is very efficient, with only three million parameters, making it feasible to run even on GPU-equipped current laptops.

We demonstrate CryoSamba’s efficacy on five cryoET datasets with three distinct voxel resolutions (achieved by pixel binning). CryoSamba substantially increases SNR for all voxel resolutions, verified both visually and quantitatively. Analysis of spatial frequencies in Fourier space confirms that our approach suppresses higher frequencies less severely than do current methods. We benchmarked CryoSamba’s performance by evaluating the Fourier Shell Correlation (FSC) (Harauz et al., 1986) for subtomogram-averaged virus-particle images, starting with tomograms before and after denoising. Comparison with the known virus structure showed that CryoSamba denoising preserved higher resolution information more faithfully than did Topaz-Denoise or CryoCARE, two widely used denoising methods.

## RESULTS

### Cryo-ET data sets

To assess the denoising capabilities of CryoSamba, we employed five distinct tomograms derived from cryo-ET of various biological samples. Two of these tomograms were obtained from the edges of plunge-frozen human BSC1 cells grown overnight on top of the electron microscopy grids, showcasing cross-sections of the plasma membrane, mitochondria, and an early endosome. These images also contained numerous free ribosomes and actin filaments within the cytosol, as well as rhesus rotavirus particles in the surrounding medium, from a study of the initial stages of virus entry (Herrmann et al., 2021; de Sautu et al., 2024).

The remaining three tomograms came from lamellae prepared by cryo-focused ion beam (cryo-FIB) milling of plunge-frozen yeast cells. These samples showed cross-sections of mitochondrial and endoplasmic reticulum (ER) membranes, ribosomes, either free in the cytosol or attached to the ER cytosolic face, and cross-sections of the double-membrane nuclear envelope. They also included some actin filaments in the cytosol.

The 3D tomographic reconstructions were produced from tilted images recorded at a nominal (unbinned) pixel size of 2.62 Å. Before denoising, we applied 3D contrast transfer function (CTF) correction using NovaCTF (Turoňová et al., 2017) to mitigate potential confounding effects from defocus. The tomograms used in our denoising tests were derived from data at various binning levels: unbinned (2.62 Å/pixel), 3x binned (7.86 Å/pixel), or 6x binned (15.72 Å/pixel), and some tests used tomograms generated from the even or odd frames of tilt series.

### Deep learning CryoSamba training pipeline

The general strategy is shown schematically in Fig. 1A. It repurposes the deep learning model Enhanced Bi-Directional Motion Estimation (EBME) (Jin et al., 2023), initially designed to enhance the frame rate of 2D videos through synthetic video frame interpolation. We treat our tomograms as “videos”, converting the z spatial direction in the tomogram into the time dimension of EBME. EBME then generates a series of “motion-compensated” xy planes. For each plane, the model generates a set of “best guesses” from pairs of planes equally spaced to either side of the xy plane in question. It then takes the average of these best guesses as the new estimate of that plane. This averaging process produces a denoising effect like that obtained by an increase in imaging exposure time (Mildenhall et al., 2018).

**Figure 1.**
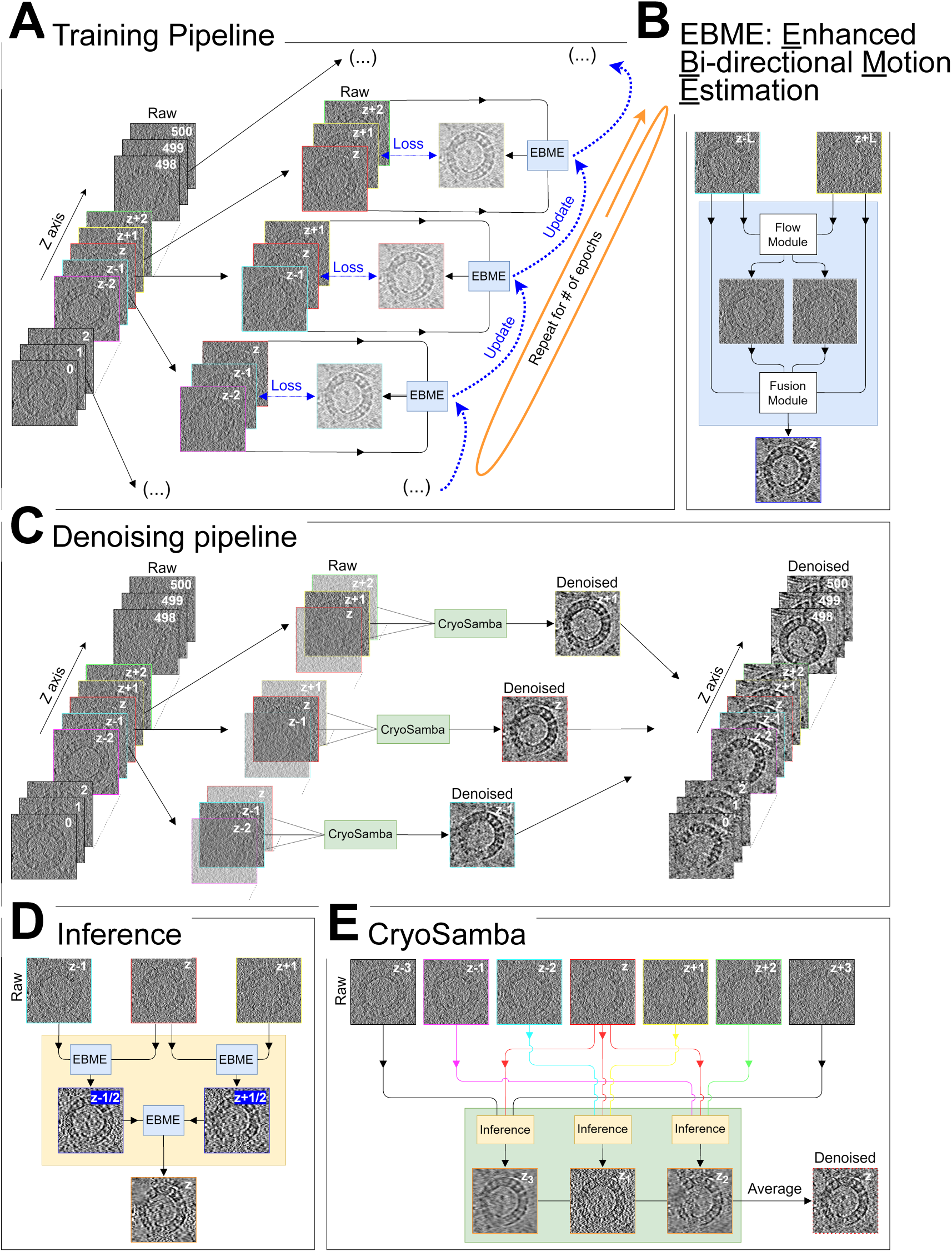
CryoSamba Pipeline for Deep-Learning Neural Network Denoising. **(A)** Training Pipeline: This component uses sets of three xy planes at z-axis positions z-L, z, and z+L. The planes at z-L and z+L are input into the EBME deep learning model. The model’s loss function evaluates the mismatch between its output and the middle plane at position z, supervising and refining the training. This iterative process is repeated for every z position (ranging from 1 to 500 in this example), with updates to the learned model weights at each step. A complete cycle (“epoch”) involves iterating through the entire dataset of images. Reaching a predetermined convergence criterion generally requires several epochs. **(B)** EBME Model: This model comprises two main modules. The Flow module assesses the bi-directional optical flow between the two input images; the Fusion module integrates the input images along with the calculated flow to predict an interpolated image. This interpolated image is positioned as if halfway between the two inputs when viewed as part of a temporal sequence. **(C)** Denoising Pipeline: Each xy plane from the tomogram is grouped with its adjacent z-1 and z+1 planes and processed through the trained CryoSamba module. This results in a denoised xy plane that supplants the original in the completed volume. **(D)** Inference Module: For each triplet, the extremal planes z-1 and z+1, along with the middle z plane, are fed separately into the EBME model. The outputs from these inputs are then reintroduced to the EBME model to refine and produce the final version of the middle z plane. **(E)** CryoSamba Module: This phase involves iterative processing, in which a specific plane z and a series of up to L_max_ surrounding planes are input into the inference module. Each triplet, z-L, z, z+L (where 1 ≤ L ≤ L_max_), is processed through the Samba module as described in (D). The results are then averaged to produce a final, denoised version of plane z.

To train the EBME model, we selected three sequential xy planes from a tomogram, spaced equally along the z-axis at positions z-L, z, and z+L (Fig. 1A). The outer planes were input into a neural network, depicted by the flow module in Fig. 1B, which computed two deformation fields to morph these planes towards the middle one. The transformed planes were merged using a U-Net (Ronneberger et al., 2015) represented by the fusion module in Fig. 1B, creating an interpolated copy of the central plane. We then minimized a loss function—reflecting the disparity between this interpolated plane and the original—by gradient descent and backpropagation (LeCun et al., 2015). Training proceeded by using plane triplets across all z values and spacings L, from L=1 up to L=Lmax, and concluded when the loss stabilized. This training process effectively reduced high-frequency noise uncorrelated across planes, such as Gaussian and shot noise; these sources of noise were further damped by averaging the interpolated images for all L spacings corresponding to the same z.

### CryoSamba denoising pipeline

The general strategy of the denoising steps is represented schematically in Fig. 1C. The inference phase of CryoSamba initiates upon completion of the training step carried out with the tomogram being denoised. For any given z in the tomogram, we input adjacent xy planes at z-L and z+L into the EBME model, to generate a denoised version at z. This process is repeated for a given z plane by varying L from 1 to Lmax and averaging them to create a final xy plane at the position z. In our tests, visual inspection was consistent with enhanced denoising. The interpolation quality decreased as L increased, leading to a tradeoff between denoising strength and blurring of fine details; an optimal image was achieved by visual ‘fine-tuning’ the value of L_max_. Fig. 1 shows an example of the effect of this process in the appearance of rotavirus-particle cross section. The series of denoised images were obtained using different L_max_ values (Fig. S1) with best chosen images obtained with L_max_ of 3 (6), 6 (12) and 10 (20) for training (inference) and voxel resolutions of 15.72Å, 7.86Å and 2.62Å, respectively.

We minimize loss of information introduced by the averaging process during the inference phase (Fig. S2) by using the “one step back, one step forward” strategy outlined in Fig. 1D. This strategy, like Samba dance steps (Guillermoprieto, 1991), involves generating synthetic xy planes at z-L/2 and z+L/2, between the experimentally determined z-L and z, and between z and z+L, respectively, which are then used as additional input during inference. By restricting use of the “one step back, one step forward” approach to the inference phase, we prevent a potential model collapse during training, that could have led for example to trivial solutions such as simply copying input frames (Reda et al., 2019).

The inference phase for a given z finishes by averaging the interpolated planes associated with it and using the average to replace the data in the original xy plane (Fig. 1E); this process is repeated for all z positions, ultimately yielding a uniformly denoised 3D volume, as schematically illustrated in Fig. 1C.

We note that in the CryoSamba denoising strategy, we train separately using each tomogram we wish to denoise. We found that the results using this approach were more reliable than those from one in which we performed a single training with many tomograms and then used the trained model to denoise naïve tomograms not included in the training pool. The fully self-supervised character of CryoSamba makes this approach possible, since it only requires for training the same 3D volume that one then wishes to denoise. In practical terms, the underlying deep learning model is relatively light (approximately three million parameters), and the training times are quite short.

To illustrate the effectiveness of CryoSamba, we visually compared tomograms before and after denoising. These tomograms were generated with 3x binning (7.86 Å/pixel) from yeast (Fig. 2A-C) and mammalian BSC-1 cells (Fig. 2D). The denoised images show enhanced SNR, estimated as describe below, in a single xy plane, approximately at the tomogram midsection, orthogonal to the electron beam direction (z). Inspection of sequential planes in the 3D volumes from yeast samples (Movies 1 and 2) confirmed improved signal-to-noise throughout.

**Figure 2.**
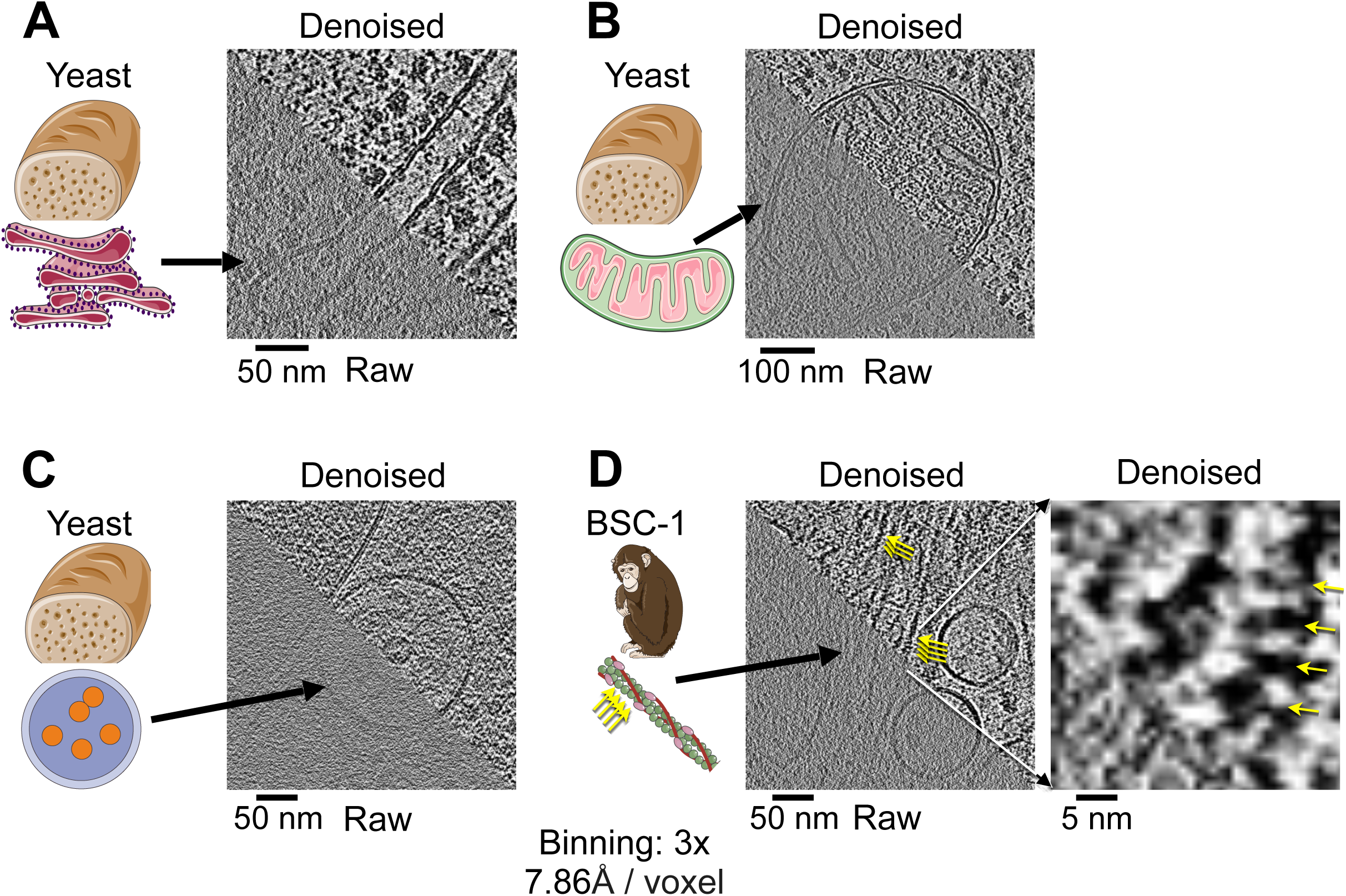
Visual Comparison of Raw and CryoSamba Denoised 3D Images of Cryo-Electron Tomograms. Visual comparison of the same xy planes from representative raw and CryoSamba denoised 3D images of cryo-tomograms, binned at 3x with a voxel resolution of 7.86 Å. **(A-C)** Different tomograms derived from three distinct yeast cells. **(A)** Cross-sections of the endoplasmic reticulum and ribosomes. **(B)** Cross-section of mitochondria. **(C)** Endosome containing intraluminal vesicles. The denoised images highlight the preservation of the double-layer appearance of the membranes, separated by approximately 4-5 nm. **(D)** Tomogram from a BSC1 cell illustrating actin filaments and the cross-section of a membrane-bound organelle of unknown identity. Distance between the arrowheads is consistent with the expected 5.5 - 6 nm periodicity of actin monomers along the helical actin filament. Scale bars: 50 nm (A-C) and 100 nm (D).

Images denoised with CryoSamba distinctly showed the characteristic phospholipid bilayer profile, a “double track” with a spacing of ∼5 nm, in membranes of the endoplasmic reticulum (ER) (Fig. 2A), mitochondria (Fig. 2B), and early endosomes (Fig. 2C, D). The enhanced clarity also resolved ribosomes in the yeast cytosol, either free (Fig. 2A, B) or interacting with the ER membrane (Fig. 2A), as well as the ∼5 nm spacing of subunits along actin filaments in the mammalian cell (Fig. 2D). Additional examples of cross sections of the plasma membrane in a BSC-1 cell and membrane-less rotaviruses in the surrounding medium are shown in (Fig. 3, panels 1-4 and panels 5-7, respectively).

**Figure 3.**
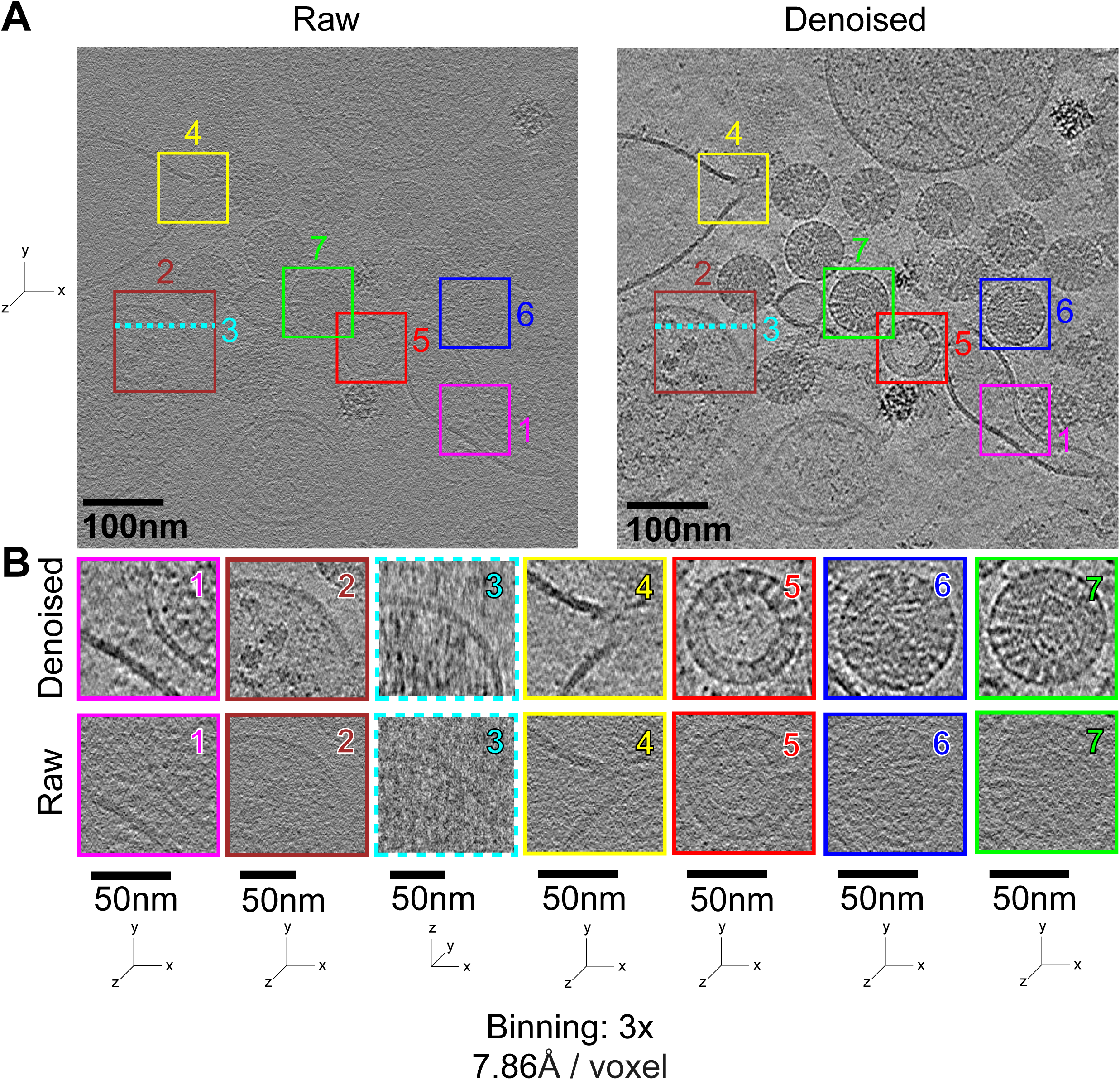
Denoising using CryoSamba of a BSC-1 Cell Incubated with Non-Enveloped Rotavirus. This figure illustrates the effects of CryoSamba denoising on a representative cryo-electron tomography image of a BSC-1 cell incubated with rotavirus. **(A)** Representative xy plane from a 3D cryo-electron tomogram obtained at 2.62 Å/voxel resolution and reconstructed to 7.86 Å/voxel resolution after 3x binning. The comparison shows the raw image before (left) and after (right) denoising. Selected regions of interest are indicated. **(B)** Enlarged xy and zx views of the selected regions before and after denoising, showing improved clarity in the images post-denoising. These enhancements are particularly noticeable in the cross-section views, showing the double-track appearance of membranes and the substructure within the virus particles. Scale bars: 100 nm (A) and 50 nm (B).

To quantify the enhancement in visual quality, we assessed the signal-to-noise ratio (SNR) determined for single viral particles in two tomograms from BSC-1 cells and throughout three tomograms from three yeast lamellae samples (Table 1). We determined SNR by two distinct methods. In one (Table 1), we generated pairs of tomograms using consecutive even and odd frames from the tilt series and calculated SNR by comparing identical xy planes in the raw data tomograms and those processed with either CryoSamba or the widely used cryo-ET denoising algorithm Topaz-denoise (Bepler et al, 2020). This method could not be used for data processed with CryoCARE (Buchholz et al, 2019), another widely used denoising algorithm, as it required consecutive even/odd tilt images for denoising; splitting them further would have degraded the CryoCARE output. To include CryoCARE in our evaluation, we used two additional approaches (Table 2): in one case, we computed the SNR using two adjacent xy planes in tomograms from the raw data and from tomograms denoised by CryoSamba, Topaz-Denoise and CryoCARE. In the second case, we estimated the SNR using the same xy plane in tomograms from the raw data and from tomograms denoised by CryoSamba, Topaz-denoise and CryoCARE. These results, summarized in Tables 1 and 2, showed comparable, substantial enhancements in SNR from both CryoSamba and Topaz-Denoise. Although CryoCARE also showed enhanced SNR, the increase came at the expense of the lower resolution imposed by the denoising algorithm (see below).

**Table 1.** SNR for images determined using alternate tilt planes and SNR ratios between raw and images denoised using CryoSamba or Topaz-Denoise. Data for two raw tomograms per experimental condition were obtained using independent sets of alternate xy tilt plane images acquired with an unbinned voxel size of 2.62 Å. SNR data for each tomogram (in dB) were calculated by correlating each xy plane throughout the volume of the two tomogram subsets (see Methods). Values in parentheses represent SNR ratios derived from the SNR values of the raw images and the corresponding SNR values of the same images denoised using CryoSamba or Topaz-Denoise (see Methods). The BSC-1 cell sample data sets capture two different rotavirus particles selected from different tomograms. The yeast sample data sets are complete tomograms obtained from three lamellae. These results, which compare raw and denoised images with the same information content, demonstrate the similar extent of SNR improvement achieved using both denoising methods.

| SNR for the images (in dB) and its ratio between raw and denoised data |  |  |  |
| --- | --- | --- | --- |
| Source | Raw | Topaz-Denoise | CryoSamba |
| Rotavirus in BSC1 cell (1) | -32.4 (0.0) | -22.9 (+9.4) | -25.7 (+6.7) |
| Rotavirus in BSC1 cell (2) | -26.3 (0.0) | -16.4 (+9.9) | -16.3 (+10.0) |
| Yeast lamella (1) | -20.2 (0.0) | -14.7 (+5.5) | -14.3 (+5.9) |
| Yeast lamella (2) | -22.9 (0.0) | -16.9 (+6.0) | -17.3 (+5.6) |
| Yeast lamella (3) | -36.2 (0.0) | -34.4 (+1.8) | -34.2 (+2.0) |
| **Total average** | **-27.6 (0.0)** | **-21.1 (+6.5)** | **-21.6 (+6.0)** |

**Table 2.** SNR for images determined at different resolutions using all tilt planes and SNR ratios between raw and images denoised using CryoSamba, Topaz-Denoise or Cryo-CARE. Data from the tomograms obtained using all xy tilt plane images of the BSC-1 cell depicted in Fig.5 acquired with voxel size of 2.62 A and processed at voxel size of 2.62 A (unbinned), 7.86 A (3x binned) and 15.72 (6x binned). SNRs were determined (see Methods) by correlating alternate xy planes (with slightly different information content) along the z axis of a given cryo-ET (Even/Odd plane) or by correlating the same xy plane (with the same information content) along the z axis determined between the raw image and the same image denoised using CryoSamba, Topaz-Denoise or Cryo-CARE (Denoise/Raw); these values are trivially infinity when the comparison is between raw and itself. The highest SNR value within a given column is highlighted in red. The results, obtained by relating raw and denoise images with the same information content highlight the extent of SNR improvement obtained using CryoSamba.

| SNR (in dB) for different voxel resolutions |  |  |  |  |  |  |  |
| --- | --- | --- | --- | --- | --- | --- | --- |
|  | Even/Odd plane SNR |  |  |  | Denoised/Raw SNR |  |  |
|  | 2.62 Å | 7.86 Å | 15.72 Å |  | 2.62 Å | 7.86 Å | 15.72 Å |
| Raw | -1.07 | 1.74 | -0.26 |  | N/A | N/A | N/A |
| CryoCARE | -1.72 | 11.98 | 17.77 |  | -28.73 | -21.83 | -22.95 |
| Topaz | 8.86 | 12.49 | 13.21 |  | 0.87 | -1.08 | 0.38 |
| CryoSamba | 7.29 | 8.10 | 8.85 |  | 3.02 | 2.64 | 6.91 |

### Comparative Impact on Resolution by Denoising with CryoSamba, CryoCARE and Topaz-Denoise

We compared the impact on resolution of denoising with CryoSamba, Topaz-Denoise and CryoCARE, by direct visual analysis of selected 2D planes and by examining the corresponding 2D Fourier transforms. As shown in Fig. 4A, while all denoising methods enhanced contrast, CryoSamba preserved high-frequency information more effectively, particularly evident by preservation of the double-track appearance of lipid bilayers in the membrane cross sections. Fig. 2 shows additional examples of membranes surrounding the endoplasmic reticulum and endosomes imaged in the yeast and BSC-1 cell samples. Inspection of the 2D Fourier transform supported our conclusions from visual inspection of the images. CryoSamba retained higher spatial frequencies that Topaz-denoise and CryoCARE flattened.

**Figure 4.**
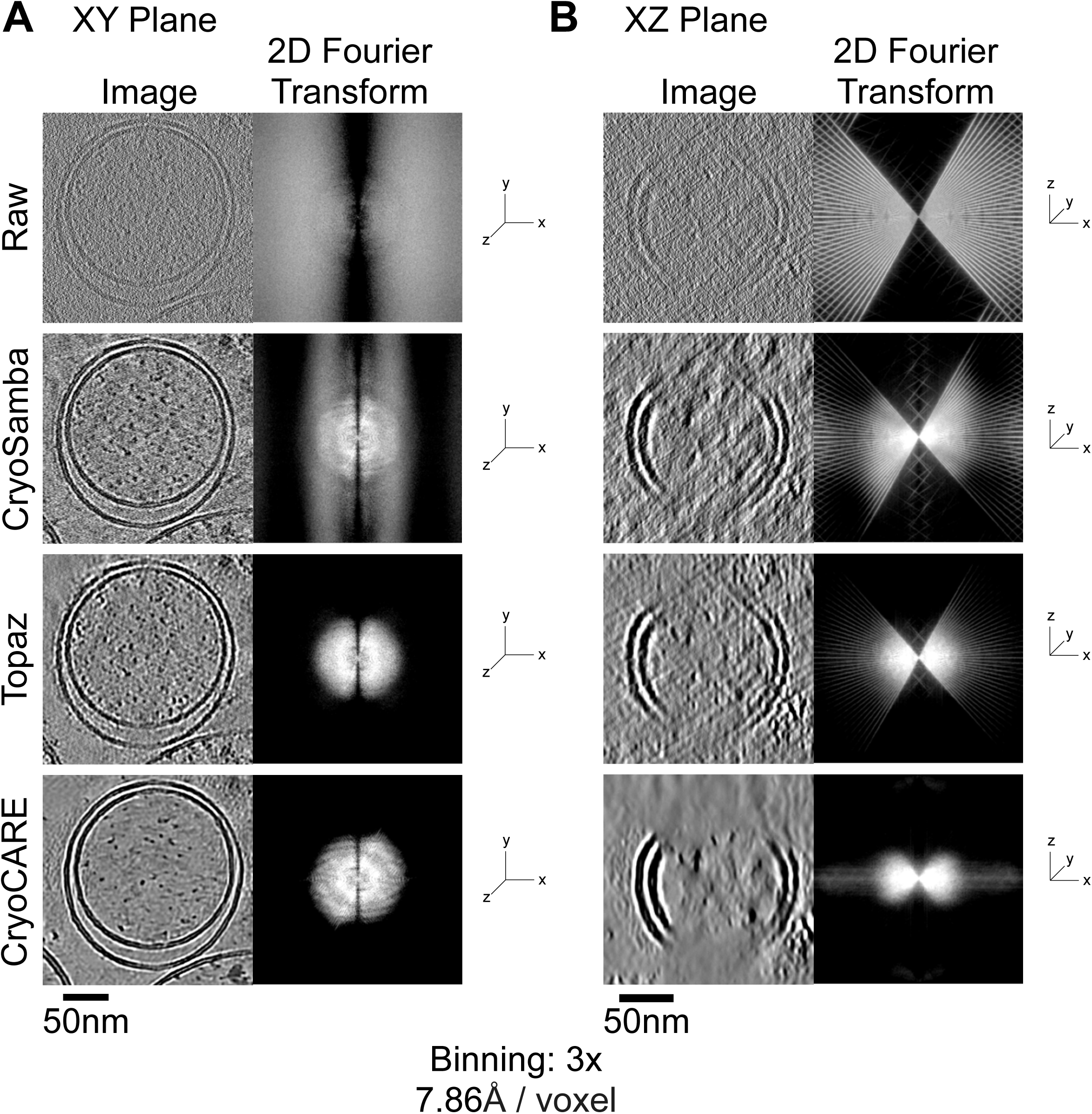
Comparison of CryoSamba, Topaz-Denoise, and CryoCARE in the 2D Fourier Plane. **(A)** Representative image from a tomogram of a BSC1 cell at 7.86 Å voxel resolution (3x binning), depicting the middle section of an organelle surrounded by two sets of membranes. The left column shows a selected xy view of a single plane of the raw image or of ones denoised using CryoSamba, Topaz-Denoise, or CryoCARE. The right column shows the (logarithm of) magnitude of the 3D Fourier transform for the corresponding regions on the left, averaged over 64 planes along the Z axis to reduce noise. The more populated higher frequency regions observed after denoising with CryoSamba are consistent with relatively better preservation of high-frequency information and the clearer double-track appearance of the membrane cross sections. **(B)** Orthogonal view of (A) corresponding to the xz plane. Inspection of the 2D Fourier transforms illustrates the extent to which high-frequency information is retained in the sample denoised using CryoSamba, the modest suppression of high spatial frequencies by Topaz-Denoise, and the dramatic loss of high spatial frequencies after denoising by CryoCARE. Scale bars: 50 nm (A, B).

We also assessed by visual inspection the denoising performance of CryoSamba, Topaz-denoise and CryoCARE for the same tomogram processed with different voxel resolutions. We selected a BSC-1 cell region containing a membrane bound organelle and rotavirus particles, and generated tomograms to final voxel resolutions of 2.62 Å, 7.86 Å, and 15.72 Å, from unbinned, 3x and 6x binning, respectively. (Fig. 5). CryoSamba yielded less blurriness, higher contrast and better-preserved double-layer appearance of the cross section of the membrane and the substructure of the virions across all resolutions, specially at 2.62 Å. Topaz-Denoise performed well at 7.86 Å, which is close to the resolution of its training data (∼10 Å), but poorly at 2.62 Å and 15.72 Å. CryoCARE had the poorest performance overall. The ability of CryoSamba to effectively handle a broad range of voxel resolutions highlights a useful versatility.

**Figure 5.**
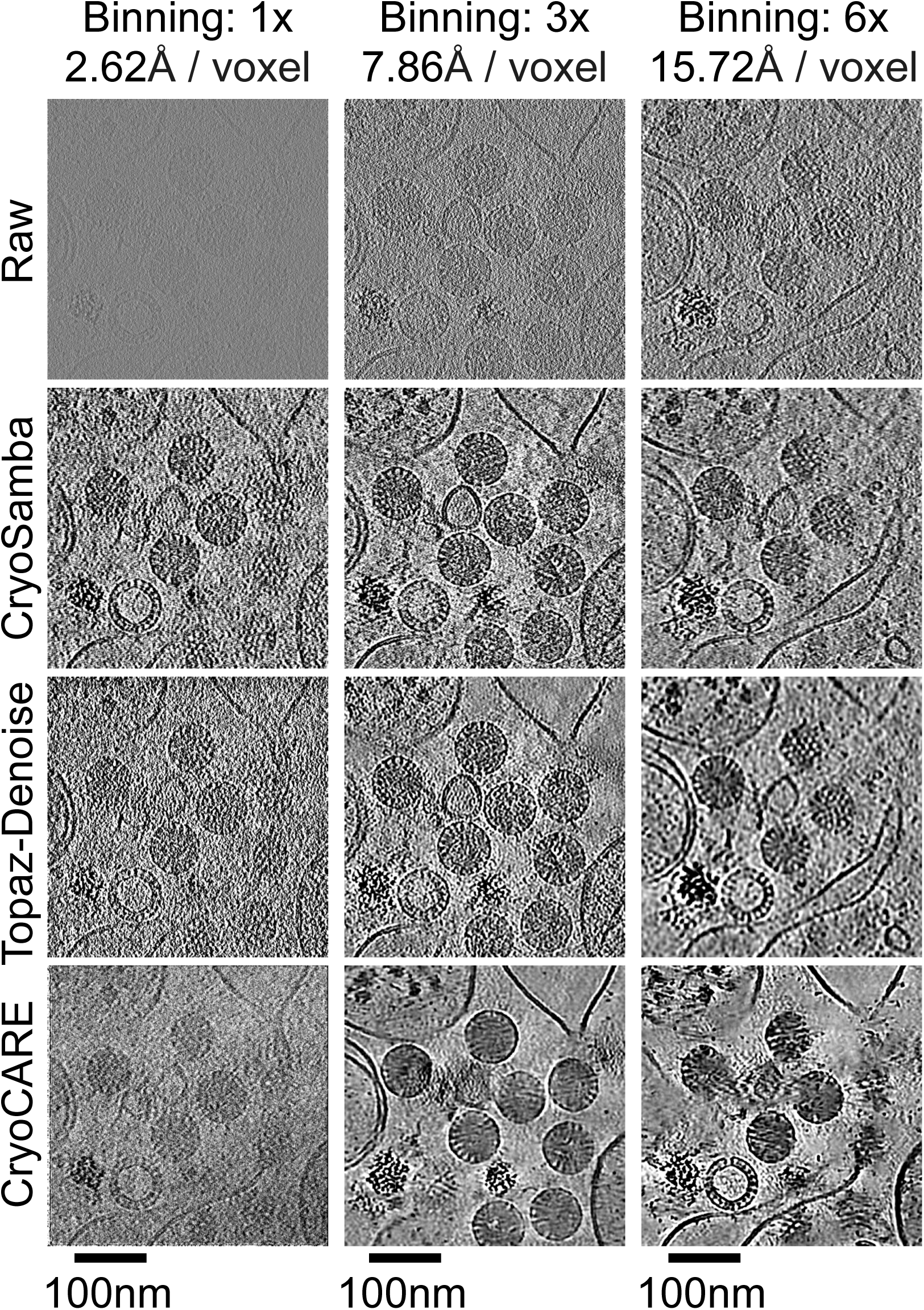
Comparison of CryoSamba, Topaz-Denoise, and CryoCARE Performance at Different Image Resolutions. Visual comparison of the same xy plane selected from the 3D reconstructed tomogram of a BSC1 cell incubated with rotaviruses, shown at voxel resolutions of 2.62 Å (left column), 7.86 Å (middle column), and 15.72 Å (right column). The images are presented before denoising (raw) and after denoising using CryoSamba, Topaz-Denoise, or CryoCARE. Scale bars: 100 nm. The images denoised with CryoSamba appear less blurred, particularly at 2.62 Å resolution and highlight the preservation of the bilayer appearance of the membrane, spaced 4-5 nm and the fine structure within the rotavirus particles.

### CryoSamba and sub-tomogram averaging

To compare CryoSamba with CryoCARE and Topaz-denoise, we used a basic STA procedure to identify and analyze rotavirus particles. We used 3D template matching and manual classification to select 54 particles across two 3x binned tomograms of BSC-1 cells (Tang et al., 2007). We carried out a single iteration of template-based alignment and averaging with final icosahedral symmetrization to achieve a simple average for feature analysis (see Methods for details). Further iterations did not improve map resolution. Since the template did not include rotavirus VP4 spikes, the presence of those spikes in the final averages indicated minimal template bias.

This analysis revealed notable differences in final map resolution. STA from tomograms derived from the raw data directly and from denoising by CryoSamba and Topaz-Denoise clearly delineated outlines of rotavirus proteins VP6 and VP7, in a T=13 icosahedral arrangement, and the projecting rotavirus VP4 spikes (Fig. 6B). The subtomogram average of the image denoised with CryoSamba also revealed VP1, the RNA polymerase surrounded by the dsRNA of the viral genome and thus in a particularly noisy environment (Fig. 6B, far right). Topaz-Denoise barely detected VP1 while CryoCARE could not.

**Figure 6.**
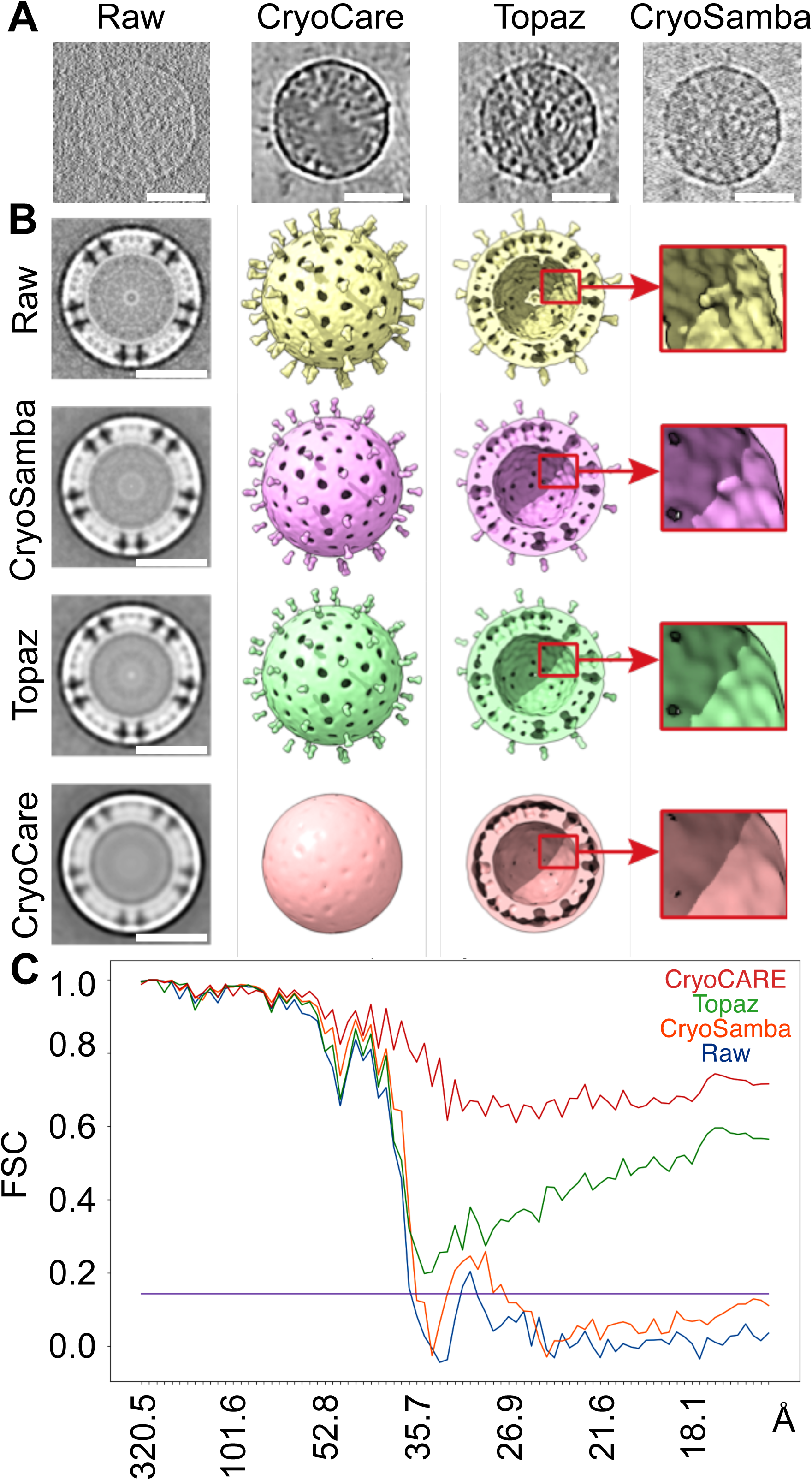
Comparison of CryoSamba, Topaz-Denoise, and CryoCARE in Information Content. **(A)** Comparison of the same xy plane selected from the 3D reconstructed tomogram of a rotavirus particle present in the sample of a BSC1 cell incubated with rotaviruses, shown at voxel resolutions of 2.62 Å (no binning) and denoised with CryoSamba, Topaz-Denoise, or CryoCARE. Spikes are visible in the particle denoised using CryoSamba and Topaz but absent in the sample denoised with CryoCARE. **Scale bar:** 35 nm. **(B)** Comparison of subtomogram averaging results obtained using raw and denoised images. The far-left column shows projected averaged maps, and the middle columns show surface renditions of the corresponding electron density maps obtained using ChimeraX. The renditions were adjusted to equivalent intensity levels of the top and middle slices. The enlarged region is centered on the location of VP1 and highlights its detection in the raw and denoised images using CryoSamba or Topaz-Denoise, but its absence in the image denoised using CryoCARE. **(C)** Comparison of the Fourier Shell Correlation calculated for the averaged subtomograms of the rotavirus before (raw) or after denoising with CryoSamba, Topaz-Denoise, or CryoCARE. The comparison highlights the similar behavior of the Fourier Shell Correlations of the raw and CryoSamba-denoised images, in contrast to the increasing values at high resolution observed for the denoised data obtained using Topaz-Denoise or CryoCARE, which appear to create high-frequency “signal” from the high-frequency noise.

We made a quantitative test of the impact of denoising on resolution by calculating a Fourier Shell Correlation (FSC) (Harauz et al., 1986) between half maps obtained by splitting the STA particles into even and odd sets and independently reconstructing separate subtomogram averages (the “gold standard” approach). CryoSamba maintained the closest correspondence to the raw data; Topaz-Denoise deviated at spatial frequencies beyond 28 Å^−1^ while CryoCARE deviated significantly at 37 Å^−1^. In other words, Topaz-Denoise and CryoCARE tend to smooth high-spatial-frequency data more severely than does CryoSamba (Fig. 6C). By preferentially enhancing contrast at the expense of losing high-resolution data, denoising with Topaz-Denoise or CryoCARE may promote self-correlation or overfitting. This potential overfitting in the Topaz-Denoise and CryoCARE data is suggested by the initial rise of their FSC curves before it reaches zero, a pattern consistent with non-independence in the masked data (Scheres et al., 2012).

## DISCUSSION

Our comparisons suggest that CryoSamba enhances both contrast and SNR without suppressing or distorting signal in the 3D tomogram, particularly at higher spatial frequencies. Its capabilities derive from its fully self-supervised algorithm, which denoises by directly engaging the 3D image itself. The optical flow interpolation method (Fig. 1B) reproduces 2D planes by reutilizing adjacent planes, not by generating new ones from scratch. This approach decreases artifacts and aids in the accurate reconstruction of high-frequency signals, which are often challenging for convolutional neural networks to capture (Rahaman et al., 2019). Moreover, the “one step back, one step forward” feature (Fig. 1C) further increases fidelity by amending residual deformations along the Z axis, effectively recycling information from the target plane to its copies.

CryoSamba circumvents the inherent limitations common to denoising techniques that rely on synthetic data (Zeng et al., 2024), which may not generalize effectively to experimental images of varying characteristics or resolutions. It also avoids the pitfalls of methods that depend on noise modeling (Li et al., 2022), which can fail to fully grasp the complexities of noise in experimental data. Moreover, CryoSamba is not contingent on the use of paired volumes (Buchholz et al., 2019; Bepler et al., 2020), which are prone to misalignments or subtle signal discrepancies.

A noteworthy advantage of CryoSamba is its ability to effectively denoise images with extremely low SNR in the raw data. For example, in Fig. 3, CryoSamba successfully denoised a tomogram processed at 2.62 Å/voxel, making apparent image details that in the raw image were almost indistinguishable from background. This capability of CryoSamba significantly broadens the direct visual inspection and potential analytical possibilities for postprocessing of data that might otherwise be considered unusable, particularly in cases involving thicker samples or data collected with reduced electron beam exposure times. The SNR and the improved contrast of the high-resolution raw image processed with CryoSamba matched those of the same volume typically associated with higher SNRs when down sampled by 3x or 6x pixel binning.

In contrast to denoising with Topaz-Denoise and CryoCARE, which in part function as low-pass filters by eliminating higher frequencies, CryoSamba better preserves most high-frequency information. This ability of CryoSamba to maintain high resolution in denoised tomograms opens new avenues for using the denoised images in post-processing tasks that would traditionally require downsampling the raw image, potentially compromising the visibility of finer details and smaller structures.

A visual comparison of averaged subtomograms of rotavirus particles added to BSC-1 cells, together with a quantitative analysis of the corresponding 3D Fourier domain spectrum from the raw and denoised data using CryoSamba, Topaz-Denoise, and CryoCARE (Fig. 6), shows that CryoSamba can reduce noise while preserving essential features of the structure. It therefore has the potential to enhance the reliability of template-based particle picking and associated automated segmentation in cellular tomography.

Finally, we have engineered CryoSamba to be relatively computationally lightweight and compatible with most commercially available workstations. It is offered in two equivalent forms: one as a command-line tool and the other as a graphical user interface (GUI) executable, enhancing ease of installation and use. Upon approval of the peer-reviewed version of this manuscript, both versions of CryoSamba, along with tomograms suitable for testing, will be freely accessible via GitHub.

## CONCLUSION

One important advantage of CryoSamba is that it operates directly on 3D reconstructed tomograms, therefore bypassing the need for accessing the original tilt series data or employing specific tomography reconstruction algorithms. Consequently, CryoSamba denoising can be incorporated at earlier stages of post-processing While this direct approach to denoising 3D images significantly broadens its applicability, it is important to note that CryoSamba does not correct for imaging defects inherent in current tomographic data acquisition protocols, such as the missing wedge.

## MATERIALS AND METHODS

### Sample Preparation

*S. cerevisiae* cells were grown overnight in YPD medium, before being diluted to 0.2 OD. One yeast strain was arrested at the metaphase-anaphase transition by degrading CDC20 using an auxin-inducible degron tag; the other strain was synchronized by α-factor for two hours, then released from G1 phase and plunge-frozen approximately 80 minutes later, such that most cells should have been in metaphase. Cells were deposited on Quantifoil R2/2 copper grids (Electron Microscopy Science) and plunge-frozen using a Leica EM-GP2 plunge (Leica).

For rotavirus samples, BSC-1 cells grown on gold grids and plunge frozen after incubation with rotavirus particles was performed as described in (Abdelhakim et al., 2014). Quantifoil gold grids R 2/2 200 mesh coated with a holey SiO_2_ film (Electron Microscopy Science) were glow discharged at 15 mA for 30 sec and then sterilized with 70% ethanol for 15 min. After washing 4 times with sterile water the grids were incubated overnight with 0.1% poly-L-lysine hydrobromide (Gibco). BSC-1 cells (at a concentration of 5×10^5^ cells/ml) were plated on the grids in DMEM (Thermo Fisher Scientific) supplemented with 10% hi-FBS (Thermo Fisher Scientific) and 1% Glutamax (Thermo Fisher Scientific) and incubated for 5 h at 37 °C and 5 % CO_2_. Cells on grids were washed 3 times with DMEM (without FBS) and then the virus previously activated with 5 μg/ml trypsin was added at a MOI of 10 and incubated at 37 °C and 5 % CO_2_ for 10 min. The grids were then blotted from the back with filter paper using the sensor blotting of the Leica Em GP2 Plunge Freezer, frozen in liquid ethane, and stored in liquid nitrogen.

### Tomogram Collection and Reconstruction

We collected five tomograms for this study. All five tomograms were collected on a Thermo Fisher Krios G3i, with a BioQuantum energy filter and a K3 direct electron detector camera. All were collected in dose-fractionation mode. Yeast tomograms were collected in counting mode at 0.15 s per frame for 1.348 s and positioning of collection was controlled by PACEtomo (Eisenstein et al., 2023). Virus tomograms were collected in counting mode at 0.15 s per frame for 1.198 s and positioning of the collection was controlled by SerialEM (Mastronarde, 2005). All collection was done at 2.620 Å/pix, or a magnification of 33k. The total dose was ∼140 e/Å^2^. Motion correction and initial alignment were done using an in-house on-the-fly pipeline, taking advantage of MotionCor2 (Zheng et al., 2017) and alignframes from IMOD (Kremer et al., 1996), respectively. Defocus files were generated by CTFFIND4 (Rohou et al., 2015).

After initial alignment, tilt series were then 3D-CTF-corrected and reconstructed using the NovaCTF pipeline (Turoňová et al., 2017), such that the whole tomogram was CTF-corrected prior to denoising and subtomogram averaging. Briefly, defocus files with a user-specified defocus step of 15 nm were applied to the CTFFIND4 files. Then, NovaCTF generated a CTF-corrected tilt series for each specified defocus step. Those CTF-corrected tilt series were individually aligned using Aretomo (Zheng et al., 2022). The resultant images were then flipped (preserving handedness of tomograms) using IMOD before filtering using the NovaCTF’s implementation in the IMOD radial filtering package. Then, tomograms were reconstructed with the NovaCTF’s 3D-CTF feature, using the relevant voxels from each CTF-corrected tilt series to reconstruct the final volume.

### Deep learning model architecture

CryoSamba uses the motion-based video frame interpolation model EBME (Jin et al., 2023), which takes as input two images (implicitly belonging to a temporal sequence) and returns a new one at an arbitrary time-point between them. As shown in Fig. 1B, this model has two main steps: bi-directional motion estimation and frame fusion.

In the motion estimation step, each input frame is initially down sampled by increasing factors of two, to form an “inverted pyramid” of images with a predetermined number of levels. At the lowest resolution level, the two corresponding downsampled images are each processed by a convolutional neural network (CNN); the correlation of the CNN output features is used to estimate two deformation fields. These fields, which can be used to “warp” one image to the other and vice-versa, are known as Optical Flow (Horn et al., 1981) and represent the (directional) pixel motion between the two frames. EBME uses a warping process known as Softmax Splatting (Niklaus et al., 2020), which directly deforms the frames to each other and combines them by a small U-Net (Ronneberger et al., 2015). At every subsequent pyramid level, the images are warped towards each other by the (upscaled) corresponding estimated flow, processed by the CNN, correlated with each other, and combined with the previous level’s outputs (as in a recurrent network (Jin et al., 2023)) to produce a refined version of the bi-directional flow. After the final level, the output flows are upscaled to the frame’s native resolution and taken to the next step.

In the fusion step, the two images are first warped towards an arbitrary time-point between them by the estimated bi-directional flow. This process assumes a linear motion model (Jin et al., 2023; Xu et al., 2019), where each flow is simply scaled by the temporal distance between the destination and its origin. In parallel, each image is also processed by a “context” CNN. The flows, warped frames, and contextual features are all combined as input to a “refinement” U-Net, which synthesizes the desired intermediate frame.

We used EBME with three pyramid levels for both training and inference, as lower values decreased performance and higher values, which increase computational costs, did not bring noticeable improvements. We also used the “high synthesis” mode (Jin et al., 2023) of EBME, which upscales flows and images by a factor of two before the fusion step, subsequently downscaling the final output to the native resolution at the end. This mode allows high-resolution processing and greatly increases the interpolation quality of small objects. The arbitrary time-point in the fusion step was always chosen to be exactly halfway between the two input images, as “asymmetric” interpolation did not work well when training the model with our chosen datasets. Finally, we used the “reflect” padding mode in all convolutional layers, to reduce the typical CNN artifacts (Alsallakh et al., 2021) at the borders of the synthesized frames, which were especially prominent after the “Samba” processes. The final model had 2.9 million weights.

### Data preprocessing

CryoSamba directly accepts as input a list of 3D image volumes in three different formats (which are automatically recognized), with no need for preprocessing and/or conversion by the user: as a single “.mrc” (or “.rec”) file, as a single “.tif” file, or as a folder containing an alphanumerically ordered sequence of 2D “.tif” images. The volumes are then initialized as memory-mapped arrays, which means that only the subregions that are currently being processed are loaded in memory, being freed afterwards. This allows us to fit very large volumes (which are typical in Cryo-ET) into systems with limited RAM.

For preprocessing, the data were converted (if necessary) to floating point values, and the intensity of each whole volume was normalized between −1 and 1. The volumes were then divided into overlapping planes of shape 256×256x1 (xyz), each of which corresponded to a square crop of a single xy plane. The crops overlapped with each other by 16 pixels along both x and y. If the number of voxels along x and/or y was not divisible by the corresponding plane size (considering the overlaps), the smaller cropped arrays at the edges were padded until they had the same shape (256×256 pixels) as the others. The padded pixels were filled not with a constant value, but with the original crop’s pixel values reflected from its border, to preserve the continuity and smoothness of the full, padded crop.

The plane crops were then combined into blocks, each consisting of three planes at the same position in x and y and at the positions z-L, z, and z+L along Z. The values of L ranged from L=1 to a fixed value L=L_max_, while z covered the whole z-axis except for the border values z<L_max_ and z>z_max_ - L_max_ that were not included. The final dataset, which consisted of the full set of blocks, was loaded into eight Nvidia A100 GPUs via Pytorch’s Distributed Data Parallel protocol (DDP) (Paszke et al., 2019; Zhao et al., 2020), in batches of 32 blocks each via 4 CPU workers.

### Training pipeline

For training, the L_max_ of the datasets was chosen as 3, 6 and 10 for pixel bins of 6x, 3x and 1x, respectively (Fig. S1). The resulting datasets were separated into training (95%) and validation (5%) splits, which were used to train the EBME model by backpropagation with the Adam optimizer (Kingma et al., 2015), with learning rate of 2e-4, a warmup scheduler for 300 iterations and a learning rate decay scheduler with a multiplicative factor of 0.99995. For each training iteration (i.e., when a single data point was passed through the model), the first and last planes of a (three-plane) block were fed into the EBME model, whose output was then compared with the block’s middle plane by photometric loss, i.e., a measure of their pixel intensity differences. The latter was a combination of a Charbonnier loss (Charbonnier et al., 1994) (a regularized version of mean squared error) with α=0.5 and ɛ=1e-3 and a ternary Census loss (a photometric loss invariant to illumination changes (Meister et al., 2018)) weighted by λ=0.1.

After every epoch, i.e., after every block of the full dataset had been passed through the model once, the block order of the training split was randomly shuffled. Furthermore, each block was transformed by randomly flipping along any of the three spatial dimensions. This transformation/deformation process, known as data augmentation (Shorten et al., 2019), increases data diversity without fundamentally altering its nature and reduces overfitting in neural networks. Training ended when both training and validation losses stabilized, which on average occurred after L_max_*30k iterations.

### Inference pipeline

For inference, L_max_ was chosen as 6, 12 and 20 for pixel bins of 6x, 3x and 1x, respectively (Fig. S1). Each dataset block was passed into a Samba module (Fig. 1D) with the underlying EBME model trained on the same data. The corresponding outputs for blocks with different inter-plane spacings L but same middle plane z were then averaged and used to compose the final volume. During inference, the latter was memory-mapped as a binary file, to fit into memory and to allow resuming the whole procedure from the last step in case it had to be interrupted. After running over the whole dataset, the final volume was converted back to its original intensity range, data type and file format.

The dataset blocks were also flipped with respect to x, y, and both x and y, and separately processed by the deep learning model. The three outputs were flipped again (undoing their transformation) and averaged with the original denoised output. This procedure (which can be optionally turned on or off in CryoSamba’s final version), known as Test-Time Augmentation (TTA) (Ayhan et al., 2018), led to a small increase in inference quality at the cost of a nearly four-fold increase in inference time.

### Hyperparameter tuning

The data block’s xy dimensions were chosen as 256×256 to include sufficient contextual information in each block, to ensure that border artifacts would not occupy a significant portion of the blocks, and to take a reasonable training/inference time without overloading the GPUs. Note that the x and y values can be chosen with a certain freedom due to the convolutional nature of the EBME model, which accepts inputs of different shapes without having to be modified or re-trained.

The loss and optimizer hyperparameters were chosen to minimize the resulting loss functions. The number of EBME pyramid levels, which is flexible enough to have different values for training and inference (Jin et al., 2023), was chosen as 3 for both routines through a grid search aimed at maximizing the final denoising performance as assessed by visual inspection.

The most important hyperparameter to tune was L_max_. For training, larger values improve the final denoising results up to a certain point, after which the improvements are negligible and increasing L_max_ only increases the computational burden (longer processing times and/or more memory usage). For inference, increasing this parameter increases the contrast enhancement effect but also the overall blurriness, so an optimized value must be found to maximize the final denoising quality. To do this, we made a simple script that, after a training run with a sufficiently large L_max_, performed inference on a small crop of the whole dataset with varying values of L_max_ and displayed the results side by side. The optimal inference value of L_max_ was then chosen by visual inspection of these results (Fig. S1), and the optimal training value was retroactively chosen (to be useful for the subsequent training runs) as half of this value. The reason for this choice is that, while training does interpolation with the z-L and z+L plane pairs of each data block, inference only does so for the corresponding (z-L, z) and (z, z+L) pairs (due to the Samba procedure), which are separated at half the distance than the former. We chose inference (training) L_max_ values of 6 (3), 12 (6) and 20 (10) for pixel bins of 6x, 3x and 1x, respectively.

### Computational requirements

The experiments reported in this work were each run on eight Nvidia A-100 Tensor Core GPUs (Table 3). Model training used about 31 GB of VRAM in each GPU and took, on average, 0.4 seconds for each iteration, with total training time (number of iterations) depending on the convergence speeds. Inference (using the trained model to denoise the data) used about 8GB of VRAM in each GPU, took about 0.9 seconds per iteration, with a completion time that was linearly proportional to the size of the volume data and the chosen L_max_. Even though training times in deep learning are usually longer than inference times, due to calculation of losses and propagation of gradients, this situation was inverted in our case due to the “Samba” and TTA procedures, which lead to 12 (3 times 4) forward passes in each iteration instead of just one.

**Table 3.** Comparison of training and inference times and VRAM usage for two workstations. The table presents a comparison of typical training and inference times, as well as VRAM usage, for two workstations employed to denoise the same tomogram using CryoSamba. The DGX-A workstation, equipped with 8 Nvidia A-100 GPUs (40GB VRAM each), utilized PyTorch’s distributed data parallel (DDP) protocol to split the training across all GPUs, and the PyTorch Compiler function to speed up computations. The LENOVO Legion Pro 5 laptop, containing a single Nvidia Geforce RTX 3070 GPU card (8GB VRAM), operated under Windows OS and could not use the PyTorch Compiler due to OS constraints. Batch sizes were adjusted to fit the available GPU memory for each workstation. Denoising calculations from a tomogram of 682 x 960 x 333 voxels, with a voxel resolution of 15.72 Å (6x binning) of a BSC-1 cell incubated with rotavirus were performed with Lmax gaps of 3 and 6 for training and inference, respectively. Times for training (per epoch) and total inference were approximately linearly proportional to the volume dimensions and maximum frame gap. Total training time is dependent on the convergence of the loss functions and tended to stabilize after around 30 epochs of training.

| Time and VRAM usage for two workstations |  |  |
| --- | --- | --- |
|  | **DGX workstation**<br>**(batch size 32, 8 GPUS)** | **Personal laptop**<br>**(batch size 4, 1 GPU)** |
| Training VRAM<br>(per GPU) | 31 GB | 6 GB |
| Training time<br>(per epoch) | 24 sec | 742 sec<br>(12 min 22 sec) |
| Training time<br>(total: 30 epochs) | 720 sec<br>(12 min) | 22260 sec<br>(6 h 11 min) |
| Inference VRAM<br>(per GPU) | 8 GB | 1 GB |
| Inference time<br>(total) | 95 sec<br>(1 min 35 sec) | 3973 sec<br>(1 h 6 min 13 sec) |

In CryoSamba, the main parameters that affect VRAM usage without significantly altering performance are batch size and number of GPUs, which defaulted to 32 and 8, respectively. Batch size defines the quantity of data passed at once to the GPUs, and running the model on N devices allows us to increase effective batch size N fold without increasing computational time. For multi-GPU runs we used DDP: the deep learning model was run independently on each device by copying all model weights for each of them and dividing the data batches among them, while using advanced procedures to synchronize their gradients during training (Zhao et al., 2020). DDP drastically improves iteration times at the cost of massively increasing VRAM usage (divided across all GPUs). We also ran the models with mixed precision (Micikevicius et al., 2017) and the optimization routines of Pytorch’s Compiler (Ansel et al., 2024) when possible (depending on the GPU’s capabilities and operating system), both of which further improved the overall computational times.

By tuning all these parameters, we could also run CryoSamba with as little as 3GB of VRAM (using batch size of 2), which fits into most personal laptop GPUs. In Table 3, we show practical VRAM usage and iteration times for a low budget laptop setup with 6 GB of VRAM and one GPU. CryoSamba is a democratic algorithm: it can run on workstations currently available to most research groups.

### Denoising using Benchmarking Algorithms

Implementation of Topaz-Denoise was done using Topaz version 0.2.5_cu11.2 (SBGrid build). The prebuilt U-Net model was used in all cases as we found it to provide superior results over self-trained models. We used the 8 GPUs available in our workstations with a patch size of 96, patch padding of 48 and no gaussian filter.

For CryoCARE, We used the CryoCARE memory efficient version (https://github.com/juglab/cryoCARE_pip) with default parameters, except for training batch size and inference number of tiles, which were chosen in order to fit the data into GPU memory. Training and inference were performed with one GPU.

### Subtomogram Averaging

Particles were selected from two rotavirus tomograms with the command e2spt_tempmatch.py in EMAN2 (Tang et al., 2007), using an atomic model of a rotavirus particle with the VP4 spikes and the interior masked out. 64 targets were selected automatically and extracted using e2spt_boxer.py. After manual inspection and removal of incorrectly selected particles, particles were aligned and averaged using e2spt_classaverage.py. Initial alignment was in c1, and icosahedral symmetry was then applied to the final map. The coordinates of particles were taken from the autogenerated “info” file by e2spt_tempmatch.py and used to extract sub volumes in each of the denoised volumes. The same subtomogram averaging protocol was applied to each denoised particle set. Subtomogram averages were visualized in ChimeraX (Pettersen et al., 2021) and sharpened to equivalent relevant levels.

### Signal-to-Noise Calculations

We estimated SNR as in Bepler et al (Bepler et al., 2020), as originally described by Frank and Al-Ali (Frank et al., 1975). The cross-correlation coefficient (CCC) was calculated between tomograms reconstructed from even tilts and tomograms denoised with CryoSamba or Topaz-Denoise reconstructed from odd tilts, and the SNR obtained from:

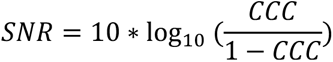

SNR was calculated for three different complete yeast tomograms and for two rotavirus particles, each selected from a different BSC-1 cell tomogram (Fig. S1, Table 1). This approach is not applicable to data denoised with CryoCARE because it requires information from all adjacent tilted frames, and hence using alternate tilts weakens its denoising capability. As a potential complementary approach, we obtained SNR values by comparing adjacent xy planes along the z axis within the raw and denoised tomograms (Table 2). We also obtained SNR values by comparing the same xy plane between the raw and denoised tomograms (Table 2). These values, however, were less informative because they correlated with the contrast enhancement regardless of preservation of the high frequency information.

---

**Figure S1.**
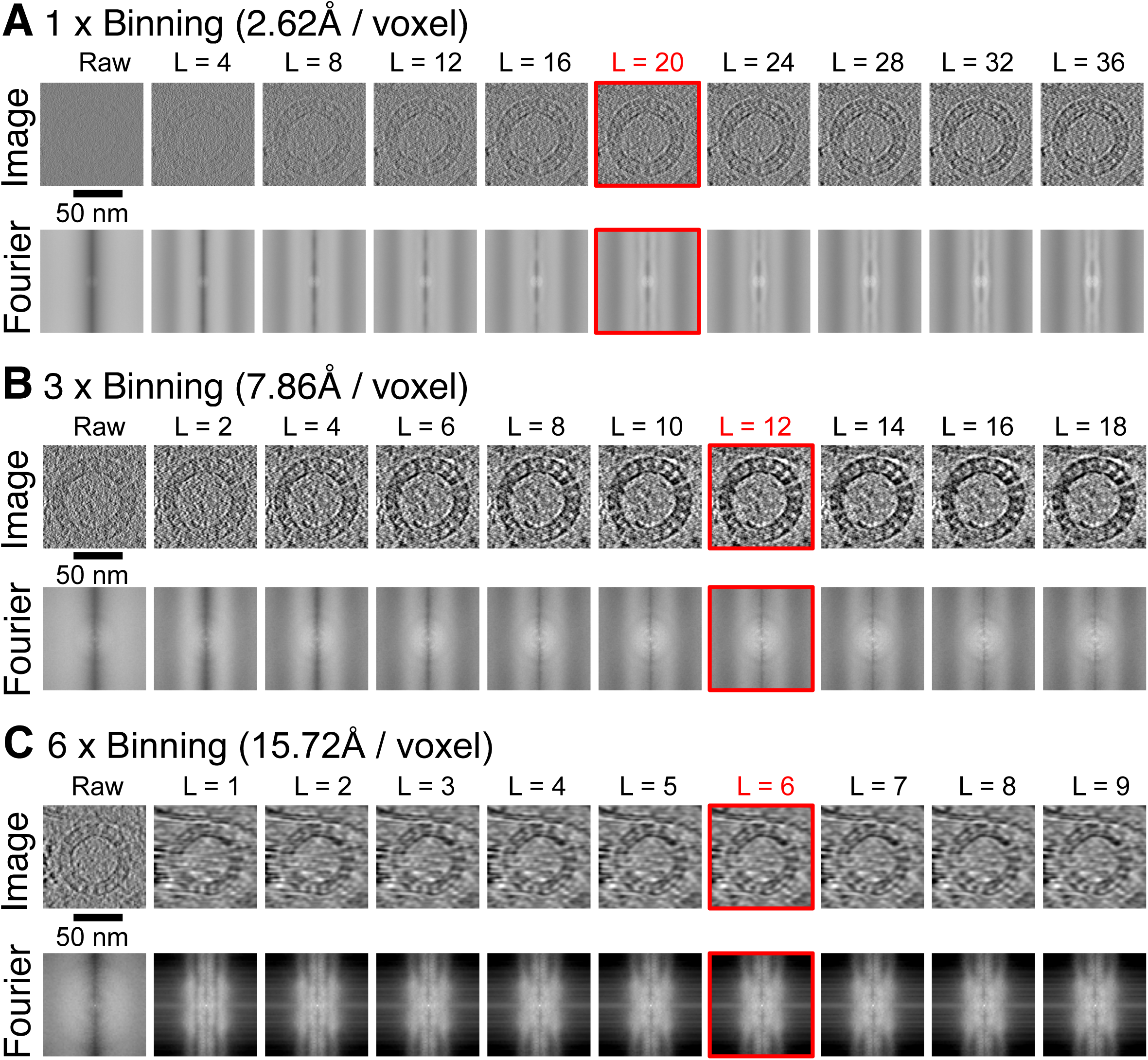
Effect of Maximum Plane Gap L_max_ on CryoSamba Denoised Images. 2D xy plane (top rows) and corresponding 2D Fourier transform magnitudes (bottom rows) of a 3D reconstructed tomogram of a BSC1 cell at voxel resolutions of (A) 15.72 Å, (B) 7.86 Å, and (C) 2.62 Å, displaying the cross-section of a defective rotavirus without spikes. Each series of images starts with the raw xy plane and is followed by the same plane after CryoSamba denoising with increasing values of the maximum plane gap hyperparameter L_max_. As L_max_ increases, both the contrast enhancement (desirable) and blurriness (undesirable) effects increase. An optimal “balancing” point (image boxed in red) is chosen based on visual inspection of these images over a reasonable range of values. The blurriness effect, which limits Lmax, is more pronounced when the relative size of structures of interest is smaller with respect to the voxel size (which dictates the “thickness” of each 2D plane), leading to smaller values of Lmax for higher pixel binning’s. Scale bars: 50 nm.

**Figure S2.**
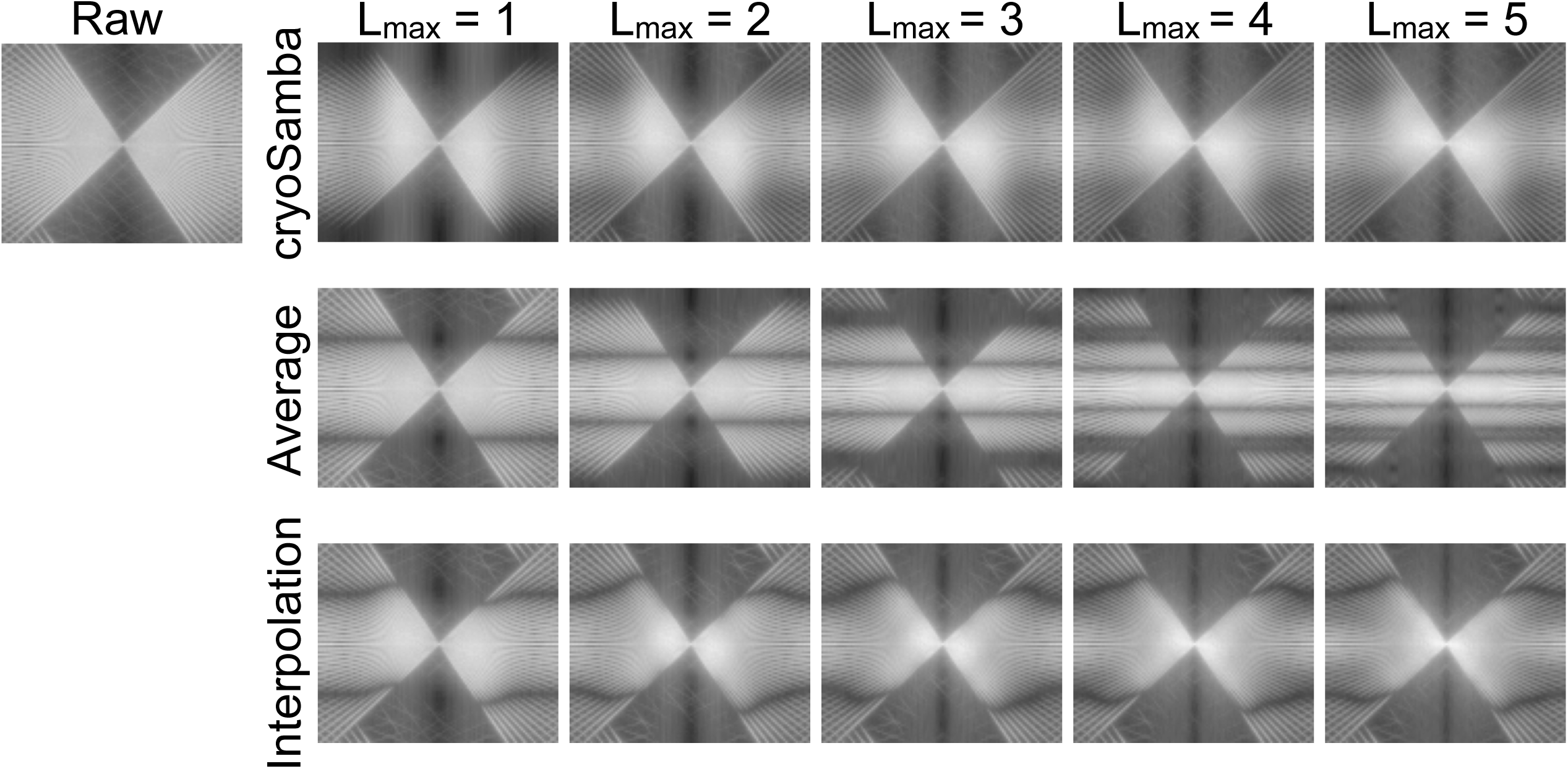
Fourier Plane Modulation Effects of Simple Averaging, Direct Interpolation, and the Samba Procedure. We apply three different denoising procedures to the xy planes of a small 3D region of a BSC1 cell cryoET tomogram and show the resulting xz Fourier transform magnitudes. The raw data image on the left displays a bundle of tilted lines characteristic of cryoET reconstructions. For the first row of data, each xy plane at its z position was directly replaced by the average of all planes between z-L_max_ and z+L_max_, for L_max_ ranging from 1 to 5. This “naïve” denoising approach enhances the imaging contrast at the cost of severe distortions, evidenced by the cosine modulations along the Z axis of the Fourier images. In the second row, the data was denoised with our trained deep learning model using a direct interpolation approach, where planes z-L and z+L are directly passed through the EBME module, and the outputs for L=1…, L_max_ are averaged. Depending on the trained model’s performance, the result could range from perfect denoising to merely averaging its inputs. The result lies between these extremes, with denoising coupled with undesirable modulation reminiscent of the averaging process. In the third row, we show the results of the CryoSamba module applied to this 3D region. The Samba procedure is designed to suppress the modulations seen in the second row. By including the original plane z in the interpolation process, we ensure that information not easily inferred from neighboring planes is propagated to the result, avoiding most of the information loss that causes the xz Fourier plane modulations. Drawings from yeast, monkey, ER and mitochondria are based on images from Servier Medical Art. Servier Medical Art by Servier is licensed under a Creative Commons Attribution 3.0 Unported License (https://creativecommons.org/licenses/by/3.0/).

## MOVIE LEGENDS

**Movie S1. CryoSamba denoising of a representative plane in a 3D tomogram from a BSC-1 cell.**

Comparison of the same image corresponding to a xy plane in a 3D tomogram volume of a BSC-1 cell incubated with rotaviruses, before and after CryoSamba denoising. The data were acquired with a per-voxel resolution of 2.62 Å and subsequently 3x pixel binned, resulting in a final resolution of 7.86 Å per voxel.

**Movie S2. CryoSamba denoising of a representative plane in a 3D tomogram from a yeast cell.**

Comparison of the same image corresponding to a xy plane in a 3D tomogram volume of a yeast cell highlighting the cross section of a portion of endoplasmic reticulum (ER) and ribosomes, free in the cytosol or associated with the ER, before and after CryoSamba denoising. The data were acquired with a per-voxel resolution of 2.62 Å.

**Movie S3. CryoSamba denoising of a 3D tomogram from a yeast cell.**

Comparison of the same tomogram volume of a yeast cell highlighting the cross section of a portion of endoplasmic reticulum (ER) and ribosomes, free in the cytosol or associated with the ER, before and after CryoSamba denoising. The data were acquired with a per-voxel resolution of 2.62 Å.

## ACKNOWLEDGMENTS

The authors thank Stephen C. Harrison for insight and editorial help, Conny Leistner for assistance in obtaining all the tomograms, Arkash Jain for assistance in the code release, Justin O’Connor for maintaining the IT infrastructure in the TK lab, and members of the TK lab for their help and support. This work was supported by a National Institute of General Medical Sciences Maximizing Investigators’ Research Award GM130386 to T.K. J.I.C.F. was supported in part by a grant from IONIS to T.K. L.M.T. acknowledges a National Science Foundation Graduate Research Fellowship under Grant DGE 2140743. M.S. was supported by a NIH Grant R01CA013202-51 to Stephen C. Harrison. The facilities of the Harvard Cryo-EM Center for Structural Biology were essential for FIB-milling the yeast cells and for recording tilt series for all the tomograms. Acquisition of the computing hardware including the DGX’s GPU-based computers, CPU clusters, fast access memory, archival servers and workstations that made possible this study were supported by generous grants from the Massachusetts Life Sciences Center to T.K. and an equipment supplement to the National Institute of General Medical Sciences Maximizing Investigators’ Research Award GM130386. Construction of the server room housing the computing hardware was made possible with generous support from the PCMM Program at Boston Children’s Hospital.

The authors declare no competing financial interests.

## AUTHOR CONTRIBUTIONS

J.I.C.F. and T.K. conceived the project. J.I.C.F. performed the denoising computational work with input from T.K. and L.T. L.T. and M.D.S prepared the samples and collected the cryoET data. L.T. calculated the subtomogram averages. J.I.C.F, L.T. and T.K. participated in data analysis and interpretation. J.I.C.F., L.T. and T. K. wrote the manuscript. All authors provided feedback and agreed on the final manuscript.

## CODE AND DATA AVAILABILITY

The code of CryoSamba and subtomogram examples used in this study are available upon request. They will be freely available in GitHub upon approval of the peer reviewed version of the manuscript.

